## Supporting information for "Facile Accelerated Specific Therapeutic (FAST) Platform to Counter Multidrug-Resistant Bacteria"

Current affiliations:

[4]Theoretical Biology & Biophysics Group, Theoretical Division, Los Alamos National Laboratory, Los Alamos, NM 87545

[5]Biotechnology & Bioengineering Department, Sandia National Laboratories, P.O. Box 969, Livermore, CA 94550

[6]Joint Bioenergy Institute, 5885 Hollis St. Emeryville, California 94608

[7]Joint Bioenergy Institute Biomass Science and Conversion Technologies, Sandia National Laboratories, P.O. Box 969, Livermore, CA 94550

[#] These authors contributed equally

### Supplementary Discussion

***Single basepair mismatch off-targets:*** α-rpsD in *S. enterica* showed homology with the start site of *rtcA* (Table S3) when allowing for zero basepair mismatches. The *rtcA* gene codes for RNA 3'-terminal phosphate cyclase and plays a role in end healing within an RNA repair pathway[1]. When allowing for a 1-bp mismatch on the start codon for all the PNAs, we found no off-targets for α-lexA, α-gyrB, α-fnr, and α-recA whereas α-ffh, α-csgD, α-folC, α-acrA, α-rpsD showed 2, 2, 3, 3, 5 off-targets respectively (Fig. S4A-D, Table S3). A single basepair-mismatch does not necessarily prevent binding to the corresponding RNA sequence but will significantly lower efficiency and increase the minimum inhibitory concentration (MIC)[2]. Using an electrophoretic mobility gel shift assay (EMSA) we find that both α-rpsD (five 1-bp mismatches) and α-lexA (zero 1-bp mismatches) show binding specificity to their target (Fig. S4E, Table S4).

***Antibiotic resistant genome screening of clinical isolates:*** CRE *E. coli* was found to have five β-lactam resistance genes, two aminoglycoside resistance genes, and one gene each for phenicol, tetracycline, sulfonamide, and trimethoprim resistance. NDM-1 KPN was found to have seven β-lactam resistance genes, three genes each for fluoroquinolone and aminoglycoside resistance and one gene each for tetracycline, sulfonamide, and trimethoprim resistance.

***Predicted RNA Folding of PNA targets:*** The folding of each PNA target theoretical sequence was analyzed using the RNAfold Web[3], looking at about 50 basepairs up and downstream of the target site. The PNA target regions of *folC* show relatively high local positional entropy, while the *rpsD*, *lexA*, and *csgD* target regions exhibit low entropy (Fig. S6). Despite this, α-folC, α-rpsD α-lexA, and α-csgD PNAs showed varying degree of effectiveness against the MDR isolates, suggesting that RNA secondary structure has little influence on PNA efficacy.

***Protein network interactions affected by knockdown PNA targets:*** Based on protein network interactions predictions from the STRING Database[4] (Fig. S7) Fnr, CsgD, and LexA tend to interact with few other proteins. Cluster coefficients indicate that these interact with very specific types of proteins that often also interact with one another. RpsD and Ffh exhibited the highest levels of nodes and average node degree. PNA targeting of *rpsD* and *ffh* are predicted to therefore result in the greatest level of cascading effects throughout dissimilar pathways in the cell and could explain the PNA's relative high success as monotherapies.

### Supplementary Methods

***Synthesis of High Concentration PNA For Toxicity Measurements:*** PNA (α-lexA and α-nonsense) used for HeLa high concentration toxicity measurements (Fig. S9) were synthesized using solid-phase chemistry on Fmoc-MBHA resin (0.22 mmol/g loading) at a 0.02 millimolar scale. Fmoc deprotection steps were performed using piperidine and coupling was performed using HATU as an activator. Post-coupling acetylation was carried out with a 5%/6% v/v solution of acetic anhydride and 2,6-lutidine, respectively, in DMF. Cleavage was performed for 2 hours using an 88%/2%/5%/5% v/v solution of TFA, TIPS, phenol, and water, respectively. The PNA molecule was precipitated in ethyl ether, centrifuged, and the purified using high pressure liquid chromatography.

***Gel Shift Mobility Assay:*** Synthetic DNA oligonucleotides that are 57-60 nucleotides in length were incubated with their respective PNA binding site at 37°C overnight. The concentration of the DNA fragments was held at 500 nM, while the PNA was always in excess at 1 μM. The reactions were performed in 1X TE with 20mM KCl (pH 7.0)[5–7]. The formation of PNA-ssDNA complexes was observed on a 20% polyacrylamide nondenaturing gel using 1X TBE running buffer. 1X SYBR-Gold® was used to stain and visualize the DNA using about 50 mL to coat the gel in low-light conditions for 20 minutes. The gel was directly put on a UV sample tray and imaged using the Gel Doc™ EZ Imaging system from Bio-Rad.

***Plasmid and strain construction:*** The SL1344-Holin strain contains a modified pRG1 plasmid that is IPTG inducible. The original pRG1 plasmid expresses the SRRz genes of the lysis cassette[8]. The lacIq gene was inserted in between the SalI and BamHI cut sites. The lacIq gene was extracted from the *E. coli* strain DH5αz1 by colony PCR with Phusion High-Fidelity DNA Polymerase (New England Biolabs) and with the following primers: forward primer (5' – AAAGGATCCCATCACTGCCCGCTTTCCAGTCG – 3') and reverse primer (5' – AAAGTCGACCCGACACCATCGAATGGTGCAAAACCTTTCG – 3'). The PCR products were subsequently gel-purified (Zymoclean Gel DNA Recovery Kit, Zymo Research Corporation), digested sequentially with *BamHI* and *SalI* (FastDigest Enzymes, Thermo Scientific) as per provided protocols, and PCR-purified (GeneJET PCR Purification Kit, Thermo Scientific) between and after digestion. The pRG1 backbone was also digested with *BamHI* and *SalI* and gel purified, and T4 DNA Ligase (Thermo Scientific) was used to ligate the pRG1 backbone and the extracted lacIq gene. Ligations were transformed into electrocompetent SL1344 cells and plasmid minipreps were performed using Zyppy Plasmid Miniprep Kit (Zymo Research). Confirmation was done by measuring the optical density with and without 1 mM IPTG (Fig. S13). The SL1344-mCherry bacterial strain contains the pFPV-mCherry plasmid (Addgene #20956) which was transformed into electrocompetent SL1344 cells and confirmed by fluorescence when excited with light at 587 nm in a light box and viewed through a 610 nm emissions filter.

***Optical Density Growth Measurements of SL1344-Holin:*** Single colonies of each bacterial strain were picked and grown in LB supplemented with 30 µg/mL Streptomycin and 100 µg/mL Ampicillin. Overnight cultures were diluted 1:1,000 into 100 µL of LB supplemented with 30 µg/mL Streptomycin and 100 µg/mL Ampicillin and 1 mM IPTG where appropriate in a 96-well plate. Cultures were grown at 37°C, with shaking, for 24 hours and $OD_{600nm}$ measured every 30 minutes using a GENios plate reader (Tecan Group Ltd.) operating under Magellan software (version 7.2).

***String Database Analysis:*** Gene names for each of the PNA-targeted RNA sequences were entered into the STRING database and searched in the organisms of *Escherichia coli* K12 MG1655, *Klebsiella pneumoniae* subspecies *pneumoniae* MGH 78578, and *Salmonella enterica* subspecies *enterica* serovar *Typhimurium* strain LT2. The meaning of network edges was set to confidence, and all active interaction sources were used except for text mining. The minimum interaction score was left at 0.400, while the max number of interactions was set to 250 for the 1$^{st}$ shell and zero for the 2$^{nd}$ shell. Counts of nodes and clusters were taken from the analysis tab.

***Osteoblast Precursor Cell Culture:*** MC3T3-E1 osteoblast precursor cells were stored in liquid nitrogen in growth media supplemented with %10 dimethyl sulfoxide (DMSO, Sigma). Osteoblasts were recovered from freezer stocks and cultured in growth media consisting of α-Minimum Essential Media (α-MEM, Gibco) supplemented with 10% Fetal Bovine Serum (FBS, Advanced, Atlanta Biologics), and 50 units/mL Penicillin-Streptomycin (P/S, Fisher Scientific) at 37°C, 5% $CO_2$, and controlled humidity. For biological replicates a freezer stock, at passage 9, was split three ways and each replicate continuously passaged as individual biological replicates at 80% confluency using 0.25% trypsin (HyClone). For toxicity experiments osteoblasts, between passages 10 and 15, were seeded on to 96-well tissue culture treated plates (Fisher Scientific) at 100,000 cells/mL in 100 µL growth media for 24 hours before infection or treatment.

***Osteoblast Precursor Infection and Toxicity Measurement:*** Osteoblast cells were infected by replacing the growth media with Dulbecco's phosphate-buffered saline containing *Salmonella* at a concentration equivalent to a multiplicity of infection of 30 and incubated at growth conditions for 45 minutes. After infection media was replaced with growth media supplemented with 30 μg/mL gentamicin instead of Penicillin-Streptomycin, incubated for 75 minutes, and replaced with fresh gentamicin containing media and treatment conditions (150 μL per well). After 18 hours of treatment 90 μL of supernatant was used to determine lactate dehydrogenase (LDH) release as a measure of cytotoxicity using the CytoSelect™ LDH Cytotoxicity Assay Kit. Percent toxicity is defined and measured as:

$$Percent\ Toxicity = \frac{\left(Abs_{450\ nm}^{Treatment} - Abs_{450\ nm}^{Negative\ Control}\right)}{\left(Abs_{450\ nm}^{Positive\ Control} - Abs_{450\ nm}^{Negative\ Contorl}\right)} \qquad \text{(Equation S1)}$$

### Supplemental Figures

| PNA | Target Sequence (5'→3') | PNA Sequence (N→C terminus) | Essentiality | Homology | | |
|---|---|---|---|---|---|---|
| | | | | *E. Coli* | KPN | STm |
| α-folC | ATACC**ATG**ATTA | $(KFF)_3$K-O-TAATCATGGTAT | Essential | **X** | | |
| α-ffh | GACA**ATG**TTTGA | $(KFF)_3$K-O-TCAAACATTGTC | Essential | **X** | **X** | **X** |
| α-lexA | CGGA**ATG**AAAGC | $(KFF)_3$K-O-GCTTTCATTCCG | Essential | **X** | **X** | **X** |
| α-gyrB | GTTG**ATG**TCGAA | $(KFF)_3$K-O-TTCGACATCAAC | Essential | **X** | **X** | **X** |
| α-rpsD | AGAAA**ATG**GCAA | $(KFF)_3$K-O-TTGCCATTTTCT | Essential | **X** | **X** | **X** |
| α-acrA | GAGGTTTACAT**A** | $(KFF)_3$K-O-TATGTAAACCTC | Non-essential | **X** | **X** | **X** |
| α-csgD | GGGGTTTCATC**A** | $(KFF)_3$K-O-TGATGAAACCCC | Non-essential | **X** | | |
| α-fnr | AGACCT**ATG**ATC | $(KFF)_3$K-O-GATCATAGGTCT | Non-essential | **X** | | |
| α-recA | **ATG**GCTATCGAC | $(KFF)_3$K-O-GTCGATAGCCAT | Non-essential | **X** | | **X** |

**Figure S1. Homology of antisense-PNA RNA-inhibitors in clinical isolates**. After sequencing, UGENE was used to search for the 12 nucleotide antisense-PNA targets in the gene of interest. The targets, predicted using the non-pathogenic and drug-sensitive strains, were present in all cases. Sequences are listed 5'to 3' with the antisense-PNA target underlined with the translation start codon in bold. Synthesized PNA sequences are listed N to C terminus starting with the CPP attached to the PNA antisense sequence by an O linker (O for AEEA). Capitol X indicates homology was found.

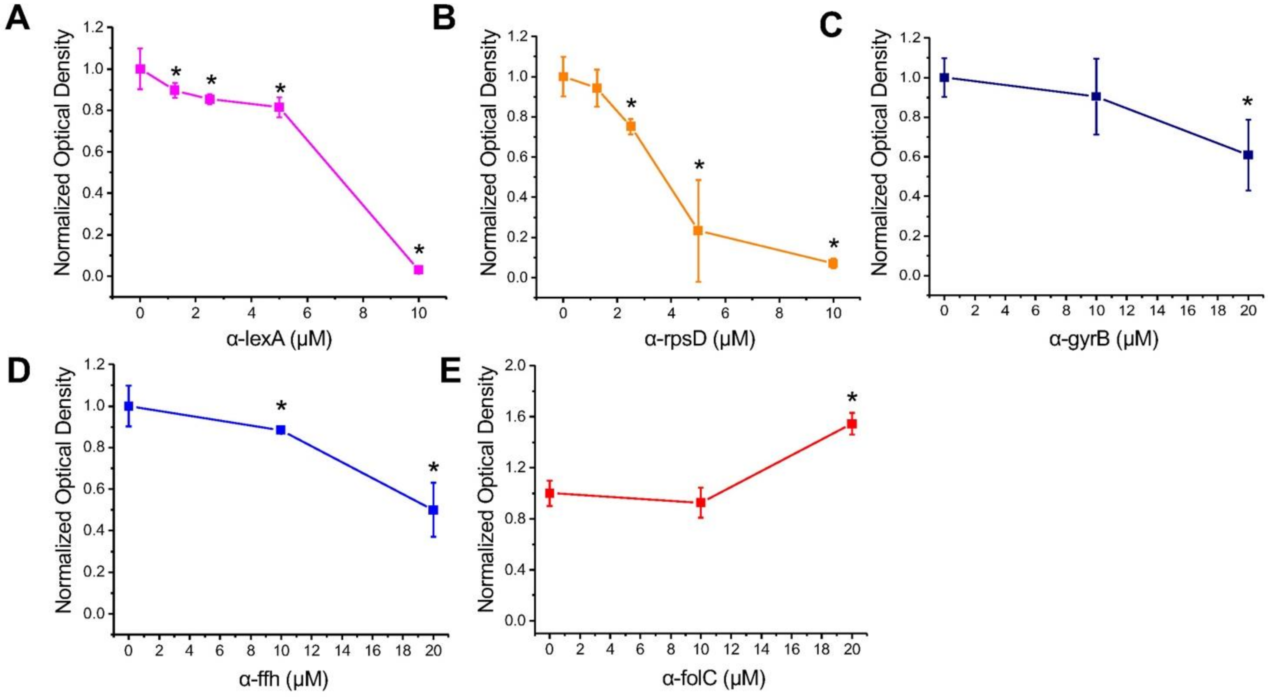


**Figure S2. MG1655 dose response of various PNA**. Optical density of a 1:10,000 dilution of MG6155 from overnight liquid culture, treated with various concentrations of antisense inhibitors was measured for 3 biological replicates with error bars as standard deviations. Normalized growth was measured as the optical density at 22 hours normalized to no treatment. An asterisk (*) indicates a significant difference ($p<0.05$) as compared to no treatment for each inhibitor.

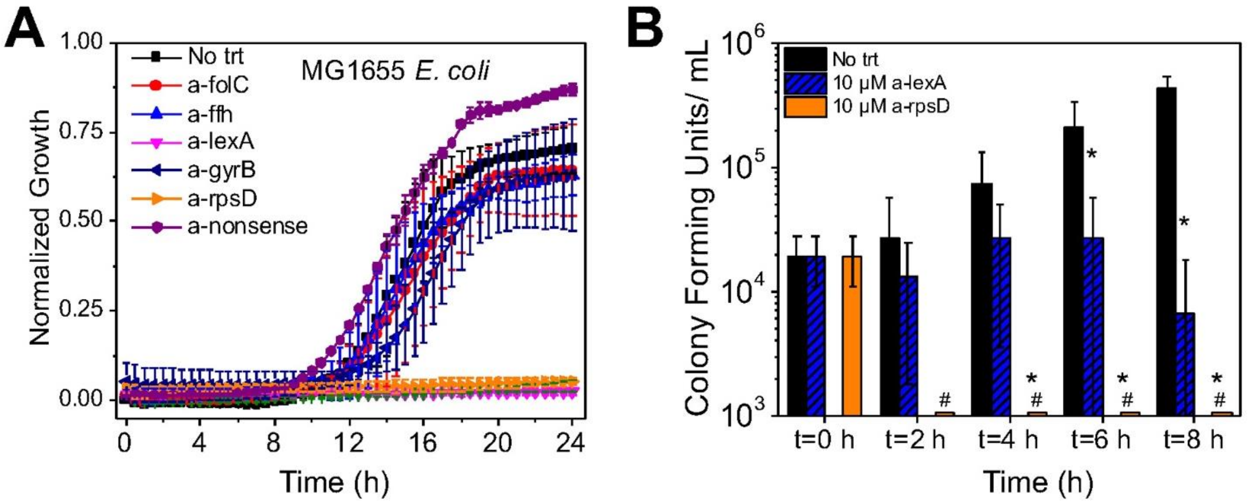


**Figure S3. Growth inhibition of *E. coli* at fixed antisense concentrations.** (**A**) Growth curves of *E. coli* MG1655 from a 1:10,000 dilution of an overnight culture, normalized to time t=0 with 10 µM of respective antisense inhibitor. Inhibitors α-lexA and α-rpsD show complete suppression of cell growth. (**B**) Colony forming units per milliliter of MG1655 *E. coli* for respective treatment as a function of time. The CFU/mL at t=0 represents the starting culture after a 1:100,000 dilution from overnight culture. MG1655 treatment with 10 µM α-rpsD resulted in 0 CFU/mL within 2 hours of treatment. Pound sign (#) represents significantly different from t=0 and asterisk (*) indicates significantly different from no treatment at each specified time point. Significance is measured as $p<0.05$ with n=3 biological replicates and error bars as standard deviations.

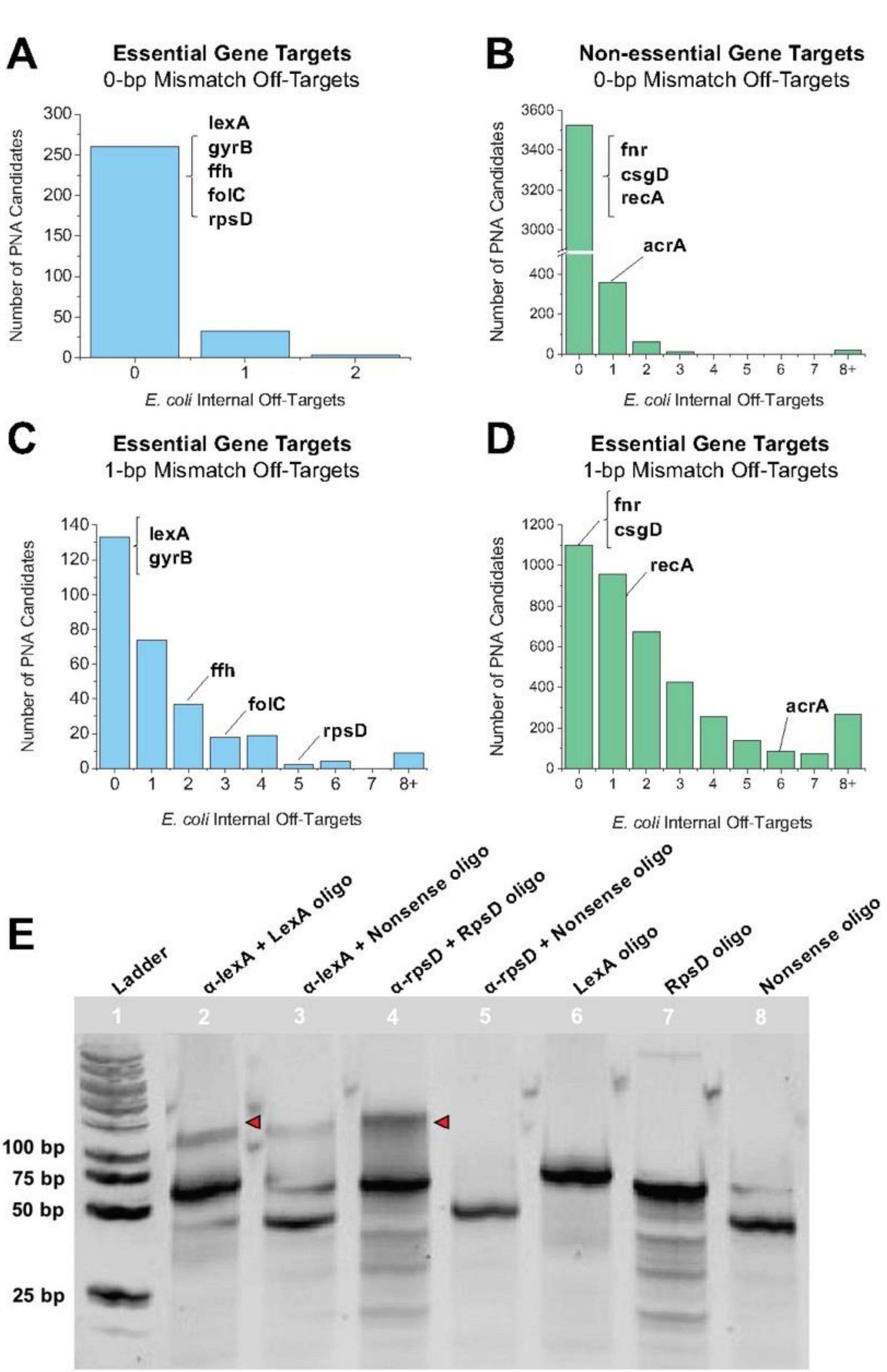


**Figure S4. PNA off-target binding.** (**A-D**) The set of potential PNA candidates were screened against the entire *E. coli* MG1655 genome to identify the distribution of off-target alignments when allowing for zero base mismatches in the alignment (A and B) and one base mismatch (C and D). In the 0-basepair mismatch analysis, all chosen PNAs that were tested had no other start codon alignments within the genome, except for *acrA* which had one, whereas in the one base mismatch alignment, the number of start codon alignments distribute across zero to six alignments. (**E**) Electrophoretic mobility shift assay (EMSA) depicting 60 nucleotide single-stranded DNA fragments containing the complementation site for their corresponding PNA. Nonsense DNA was a random 60 nucleotide single-stranded DNA sequence that contained no complementation sites for PNA binding (Table S4). Lanes with only PNA showed no bands because the SYBR-Gold Ⓡ stain intercalates between the base pairs and the PNA has no secondary structure (bands not shown). Bands corresponding to PNA + complementary DNA are shifted above the unbound DNA (red triangles). Corresponding PNA + nonsense DNA bands are seen in lanes 3 and 5, with lane 3 having slight cross-over bands coming from lane 2. All lanes are from the same gel.

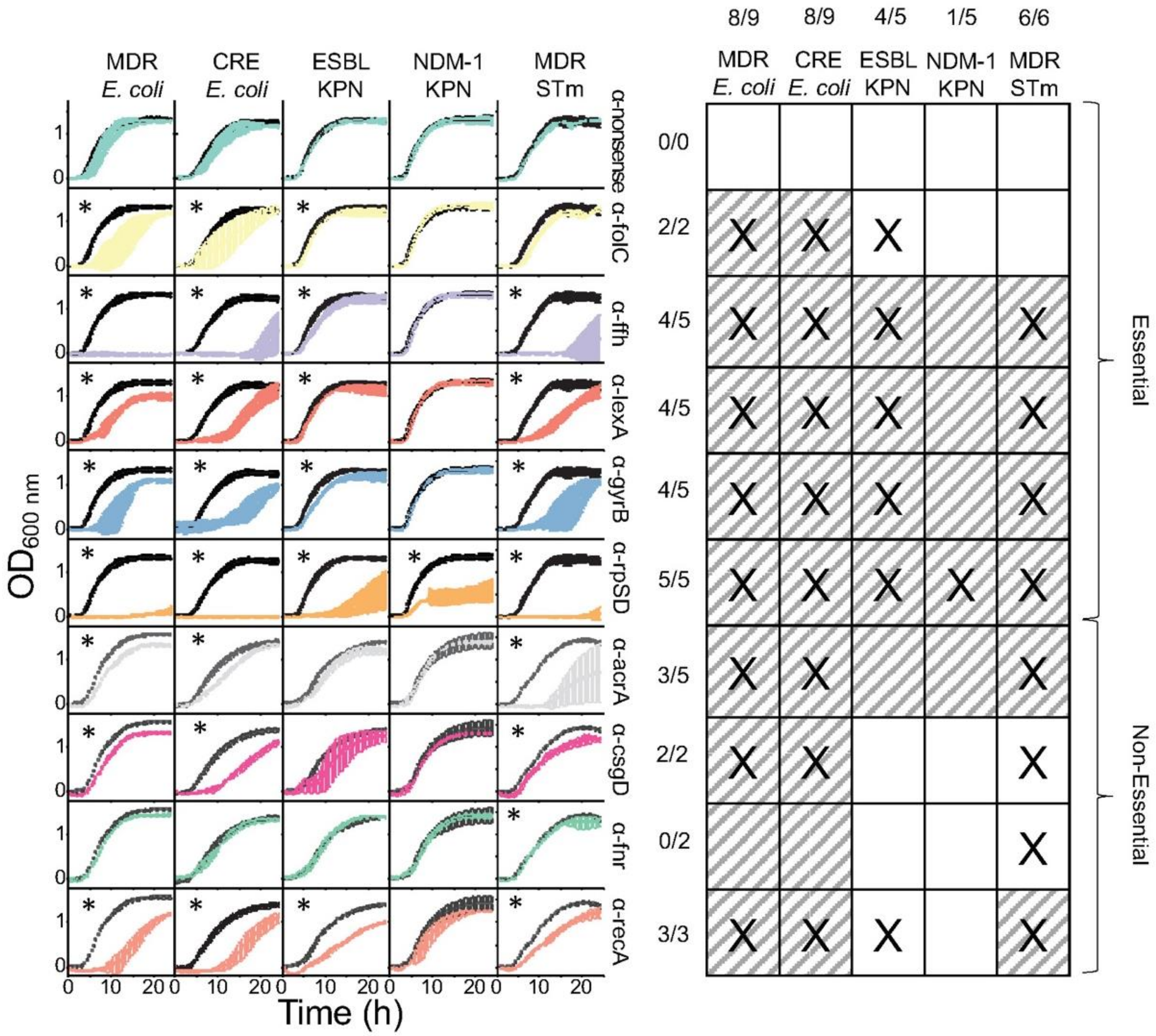


**Figure S5. Growth curves of MDR clinical isolates with respective PNA monotherapy treatment.** Growth curves shown are the average of at least three biological replicates normalized to the optical density at t=0 with error bars as standard deviation. Each antisense inhibitor treatment (10 µM) shown with no treatment growth curves in black for comparison. Data were used in Fig. 2E-F of the main text. An asterisk (*) indicates a significant difference ($p<0.05$) at 16 hours compared to no treatment at 16 hours. The table on the right indicates homology and efficacy of each PNA treatment. Grey diagonal lines indicate homology of the clinical isolate and PNA and an X indicates significant growth inhibition compared to no treatment. Fractions indicate the number of treatments that have homology with the isolate and which show a significant difference to no treatment over the number of treatments that have homology, with fractions on the left corresponding to the PNA of that row and fractions above corresponding to the indicated clinical isolate.

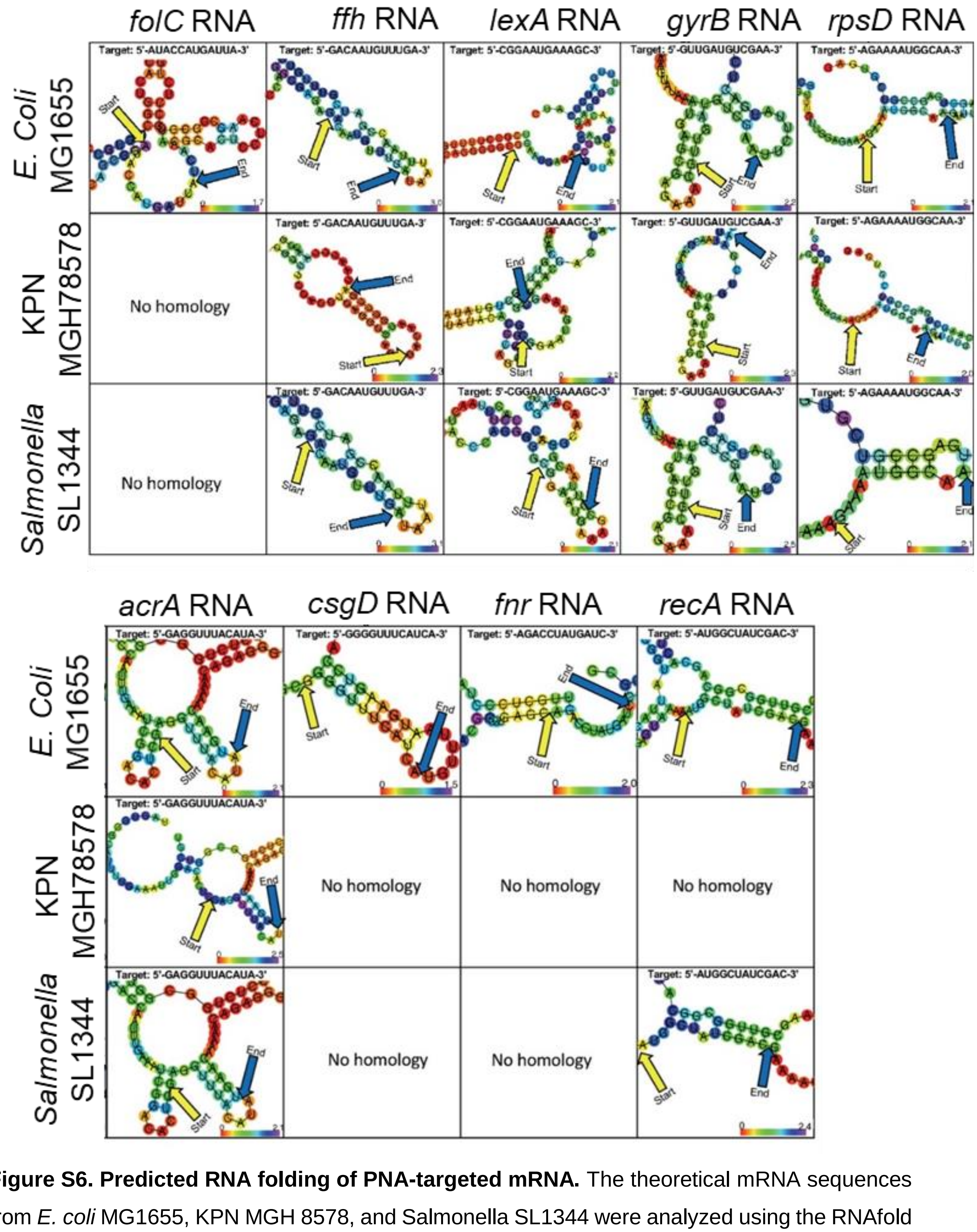


**Figure S6. Predicted RNA folding of PNA-targeted mRNA.** The theoretical mRNA sequences from *E. coli* MG1655, KPN MGH 8578, and Salmonella SL1344 were analyzed using the RNAfold

WebServer[3]. Folding is based on minimum free energy structures. Red shades indicate lower entropy, or areas of low conformational flexibility. The specific binding site of the PNA sequence is shown above. Top and bottom panel show essential and non-essential genes targeted in this study.

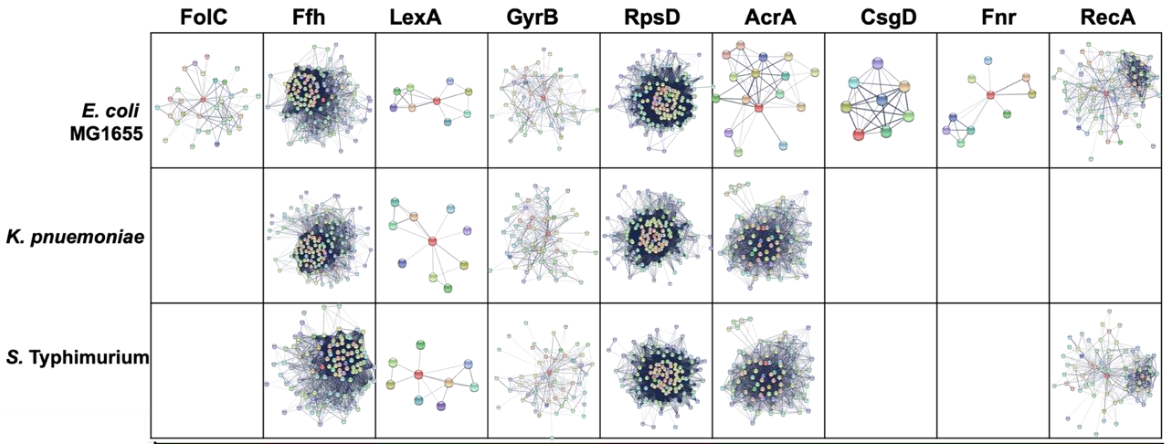


| | | FolC | Ffh | LexA | GyrB | RpsD | AcrA | CsgD | Fnr | RecA |
|---|---|---|---|---|---|---|---|---|---|---|
| Number of Nodes | *E. coli* MG1655 | 32 | 125 | 9 | 51 | 136 | 16 | 8 | 10 | 69 |
| | *K. pnuemoniae* | | 139 | 10 | 56 | 140 | 82 | | | |
| | *S.* Typhimurium | | 111 | 9 | 55 | 124 | 56 | | | 68 |
| Number of Edges | *E. coli* MG1655 | 110 | 3311 | 17 | 256 | 5042 | 45 | 28 | 16 | 509 |
| | *K. pnuemoniae* | | 3637 | 14 | 317 | 4866 | 1737 | | | |
| | *S.* Typhimurium | | 2724 | 14 | 253 | 43200 | 775 | | | 480 |
| Average Node Degree | *E. coli* MG1655 | 6.88 | 53 | 3.78 | 10 | 74.1 | 5.62 | 7 | 3.2 | 14.8 |
| | *K. pnuemoniae* | | 52.3 | 2.8 | 11.3 | 69.5 | 42.4 | | | |
| | *S.* Typhimurium | | 49.1 | 3.11 | 9.2 | 69.7 | 27.7 | | | 14.1 |
| Average Local Clustering Coefficient | *E. coli* MG1655 | 0.748 | 0.755 | 0.851 | 0.659 | 0.822 | 0.652 | 1 | 0.919 | 0.652 |
| | *K. pnuemoniae* | | 0.755 | 0.881 | 0.598 | 0.81 | 0.773 | | | |
| | *S.* Typhimurium | | 0.773 | 0.857 | 0.628 | 0.84 | 0.815 | | | 0.666 |

**Figure S7. Predicted protein interaction networks of each PNA gene target.** Known protein-protein interactions (obtained from the STRING database) between the gene targeted by each PNA and other proteins throughout the cell are shown as string maps[4]. String maps present connections based on confidence, with darker lines indicating greater confidence in the existence of an interaction and node color indicating “closeness” to the target protein. The clustering coefficient representing the “tightness” of the proteins in the network. Networks were constructed for standard genomes of *E. coli* MG1655, *K. pneumoniae*, and *S.* Typhimurium. Blanks indicated where the corresponding protein targets by the PNA has no homology in the bacteria. Networks were constructed using information from the String database based on co-expression, co-occurrence, gene fusion, experimental validation, databases, and neighborhood. A medium confidence interaction score of 0.400 threshold was established and the maximum number of networks for the first shell set to 250 and the second shell set to none. Below the respective protein network interactions is a table containing various protein interaction network values such as the number of nodes, number of edges, average node degree, and average local clustering coefficients. Each value is color coded with the lowest value of the appropriate protein network interaction value in green and the highest in red.

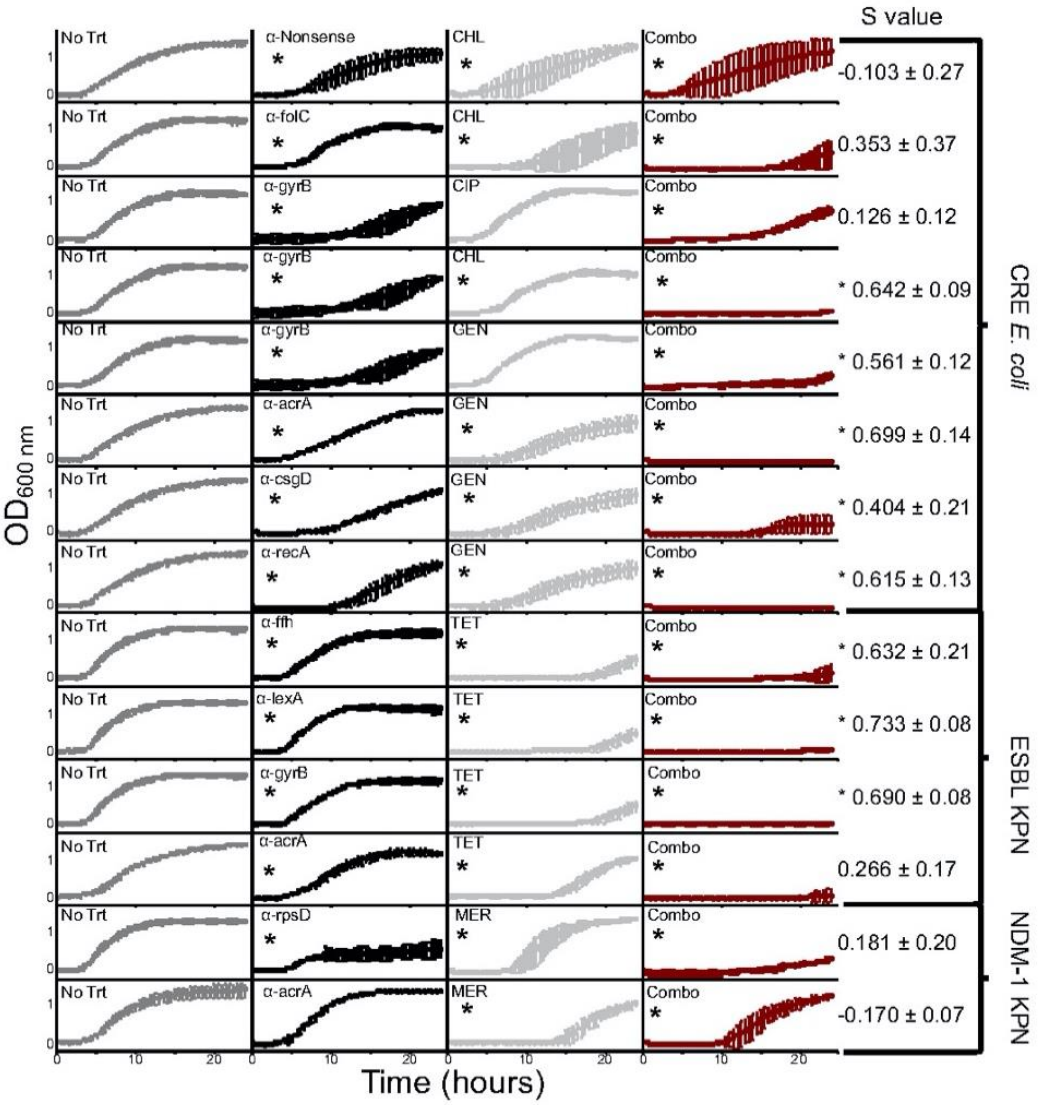


**Figure S8. Growth curves of MDR clinical isolates subjected to antibiotic and PNA combination treatment.** Optical density growth curves of each multidrug resistant bacteria from a 1:10,000 dilution of overnight liquid cultures, normalized to t=0. Growth curves shown are of at least three biological replicates with error bars representing standard deviation; the data shown was used in Fig. 3 of the main text, and an asterisk (*) indicates significant inhibition ($p<0.05$) at 16 hours as compared to no treatment. All conditions were treated with 10 µM of the indicated PNA. Antibiotic concentrations were (from top to bottom) 8 µg/mL chloramphenicol (CHL), 1 µg/mL ciprofloxacin (CIP), 4 µg/mL gentamicin (GEN), 2 µg/mL tetracycline (TET), and 8 µg/mL meropenem (MER). CHL, CIP, and GEN are all at concentrations corresponding to their CLSI "sensitive" breakpoints, TET is at a concentration below its CLSI "sensitive" breakpoint, and MER is at a concentration about its CLSI "resistant" breakpoint.

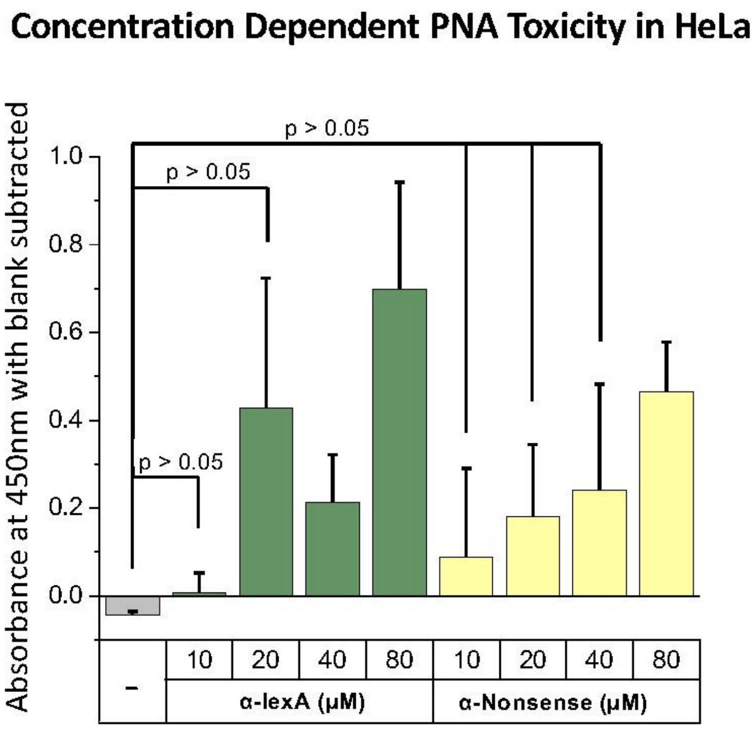


**Figure S9. Cytotoxicity measurement of varying concentration of PNA in HeLa cells** HeLa cells were plated on tissue culture treated 96-well plates at 4,500 cells per well. Varying concentrations of α-lexA and α-nonsense were added exogenously after 24 hours of growth and incubated in growth conditions for 18 hours. Cytotoxicity was measured using a lactate dehydrogenase assay and absorbance was read at 450 nm with a blank (growth media with lactate dehydrogenase assay) subtracted. All conditions were compared to the negative control indicated by a minus sign (-) and no significant difference, indicated by $p>0.05$, was seen for α-lexA up to 20 μM and α-nonsense up to 40 μM. Error bars are standard deviations.

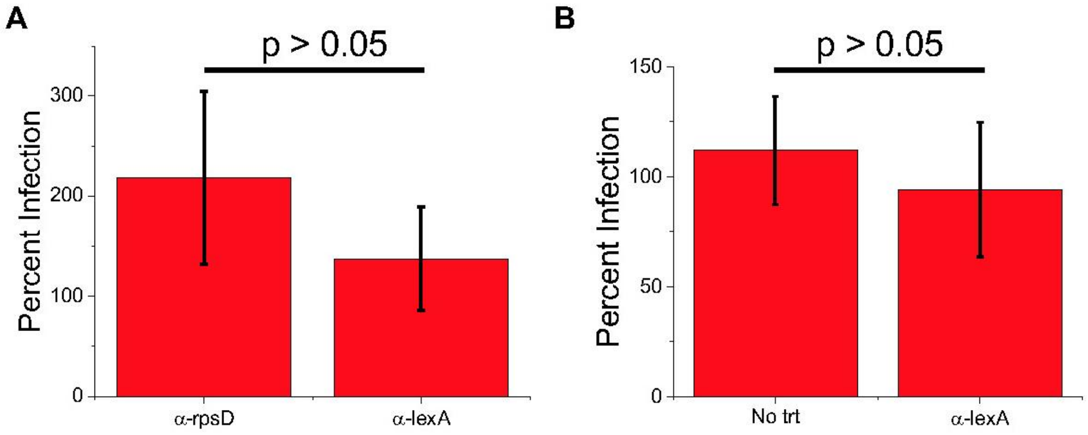


**Figure S10. Effect of 45-minute incubation with PNA treatment during infection.** (**A**) Individual colonies of SL1344 were picked off of solid media and grown for 16 hours in LB medium supplemented with 30 µg/mL streptomycin, diluted 1:10, and regrown for 4 hours. After regrowth bacteria was diluted to a concentration equivalent to a 30 MOI of 200,000 cells/mL in 100 µL (conditions of HeLa cell infection after 24 hours growth). Cultures were suspended in phosphate buffered saline and treated with 10 µM of either α-lexA, α-rpsD, or PSB for 45 minutes before serial dilution, plated on solid media (40 µg/mL streptomycin), and incubated for 16 hours. Colony forming units were counted to enumerate percent infection with respect to no treatment. There is no significant difference ($p>0.05$) between α-lexA and α-rpsD at 45 minutes of treatment. (**B**) Cultures were grown and treated as described for panel A except that prior to treatment an aliquot was removed from each biological replicate as a no treatment condition for that biological replicate. Percent infection was then calculated in comparison to each replicates' no treatment. Error bars are standard deviation for three biological replicates.

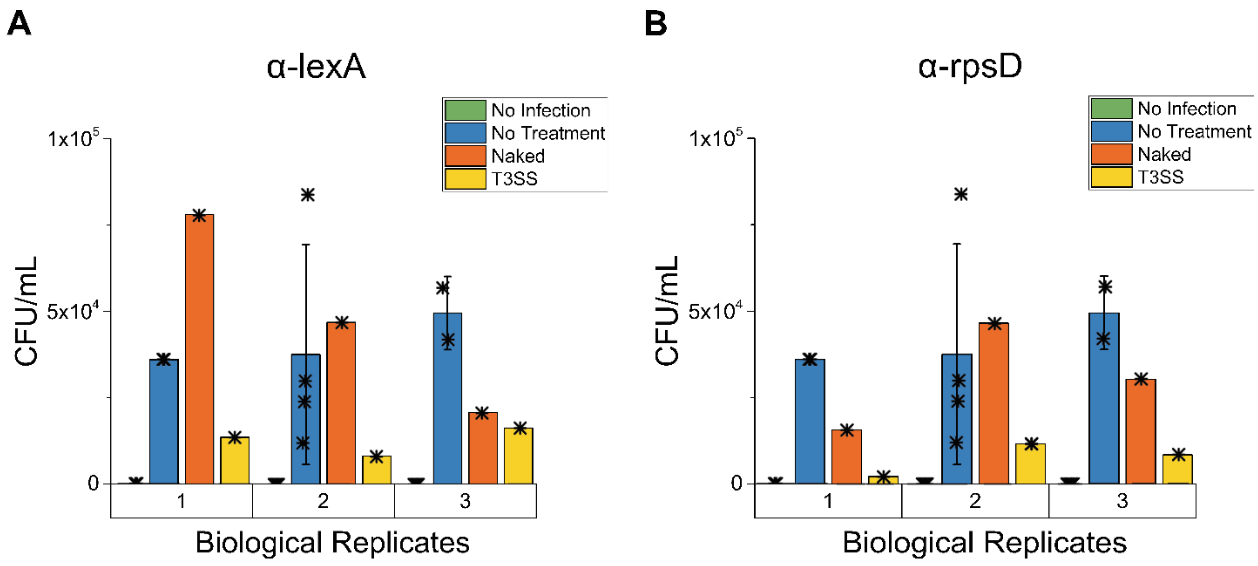


**Figure S11. Raw CFU/mL of HeLa infection treated by T3SS.** Intracellular STm-GFP (SL1344) colonies were measured by lysing infection HeLa cells with 30 µL of 0.1% Triton for 15 minutes at room temperature, followed by 1:10 dilution by addition of 270 µL DPBS. The lysate was serially diluted 10X, and 10 µL were plated onto LB agar with 40 µg/mL of Streptomycin. Following overnight growth colonies were counted to determine the CFU/mL. Shown are individual HeLa biological replicates shown in Figure. 4C where error bars are standard deviation between technical replicates shown as asterisks.

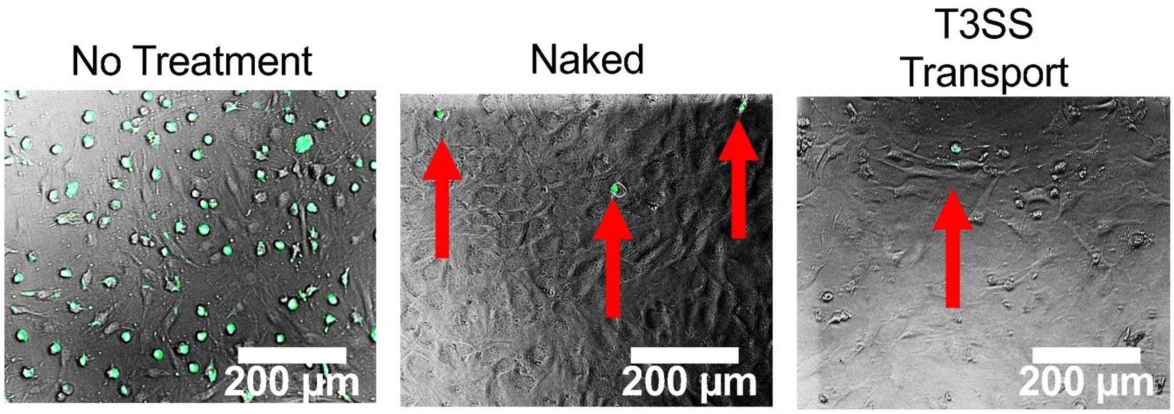


**Figure S12**. **T3SS-PNA treatment eliminates *Salmonella* infection of osteoblast cells**. Osteoblast precursor cells were grown for 24 hours on tissue culture treated 96-well plates prior to infection with STm- GFP (SL1344) at a multiplicity of infection of 30. The antisense inhibitor α-rpsD was added at 10 µM either without infection (No Treatment), after infection (Naked), or during infection (T3SS-PNA). After treatment for 18 hours the infection was imaged in brightfield and GFP channel. GFP expression corresponds to an intracellular infection with red arrows indicating hard to see GFP expression.

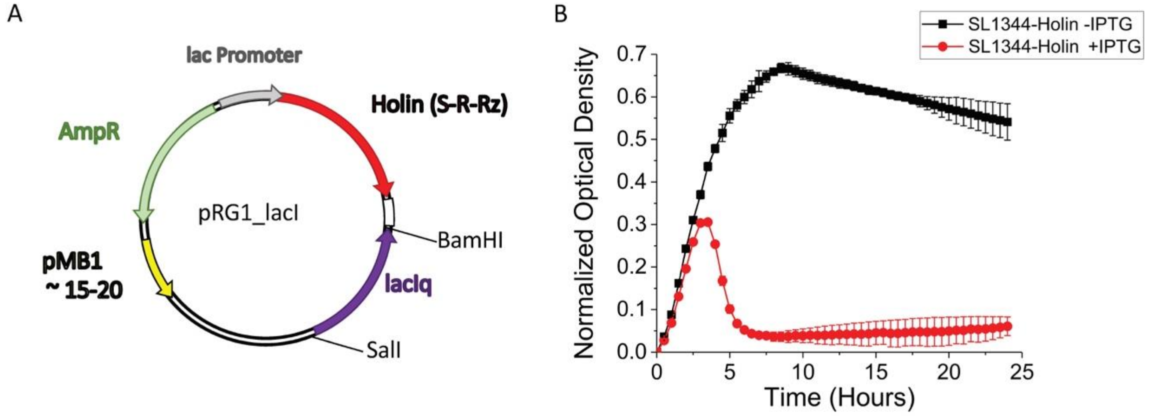


**Figure S13. Plasmid map and growth curves of IPTG induced lysis of SL1344-Holin.** (**A**) Plasmid map of pRG1 modified to include the lacIq gene showing the Holin-Endolysin cassette, Ampicillin resistance gene, lac promoter, and pMB1 origin of replication. (**B**) Optical density of SL1344-Holin (Delivery STm) with or without 1 mM IPTG was measured at 600 nm and normalized optical density is reported as absorbance at 600 nm minus media blank and each biological replicate normalized to the optical density at time zero. 1 mM IPTG was added for induction. Error bars indicate standard deviation of n=3 biological replicates.

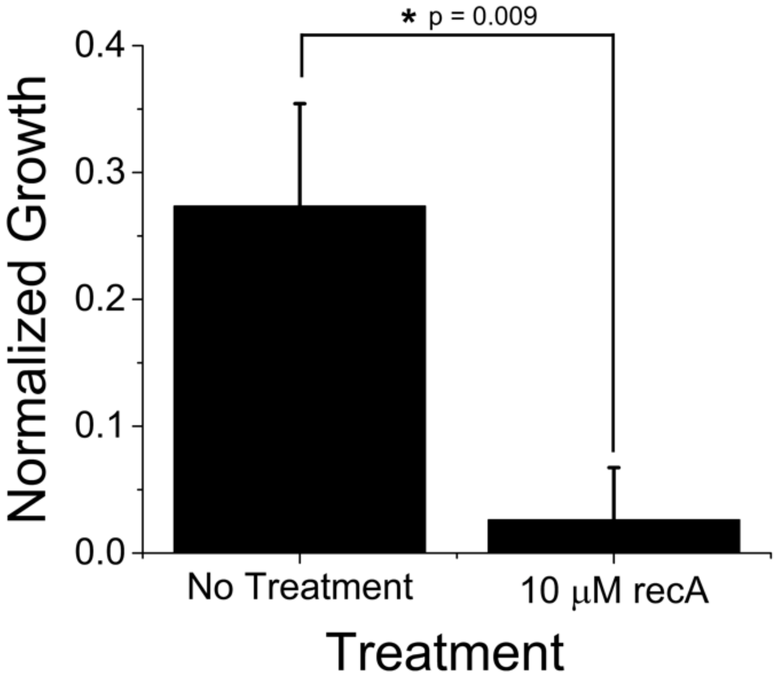


**Figure S14. α-recA growth inhibition of SL1344.** Optical density was measured at 600 nm in 50 µL cultures and normalized optical density is reported at 24 hours as absorbance at 600 nm minus media blank and each biological replicate normalized to the optical density reading at time zero. Overnight cultures of SL1344 were diluted 1:10,000 and treated with10 µM addition of α-recA for 24 hours. Error bars indicate standard deviation of n=3 biological replicates.

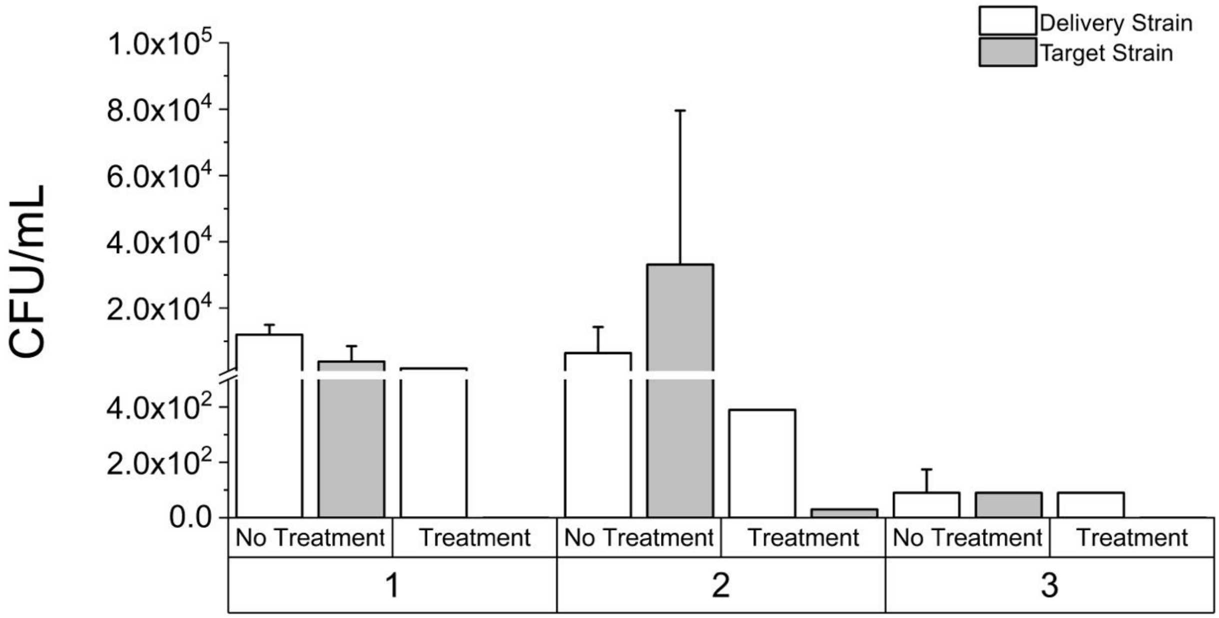


**Figure S15. Raw CFU/mL in double infection study.** Following double infection experiments colony forming units were counted for both Target STm (SL1344-mCherry, determined by florescence) and Delivery STm (SL1344-Holin) (Fig. 4E). Intracellular SL1344 colonies were enumerated by lysing with 30 µL of 0.1% Triton for 15 minutes at room temperature then diluted 1:10 by addition of 270 µL of DPBS. The lysate was serially diluted 1:10 in 100 µLs and 10 µL plated onto LB agar with 40 µg/mL of Streptomycin and 100 µg/mL Ampicillin. Error bars indicate standard deviation of technical replicates. The biological replicates are indicated by number 1,2, and 3 on x-axis.

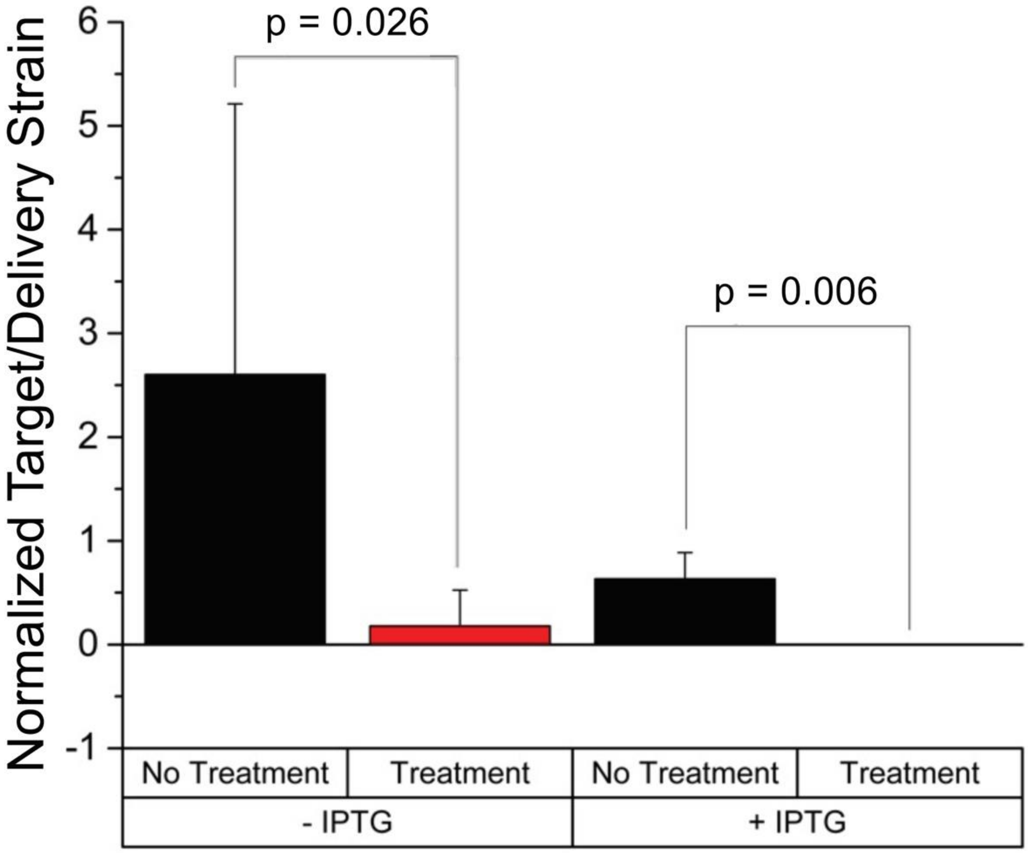


**Figure S16. Normalized Target STm to Delivery STm treatment with IPTG induction of PLac promoter.** The double infection clearance described in Fig. 4D but in presence of 1 mM IPTG to induce pLac promoter controlling expression Holin/Endolysin gene in Delivery STm to release PNA (α-recA). Significant decrease in the normalized Target STm to Delivery STm was measured in both absence (-IPTG, p=0.026) and presence of IPTG (+IPTG, p=0.006). This data showed that the leaky expression (-IPTG) was sufficient to turn on Holin/Endolysin based kill switch.

### Supplementary Tables

**Table S1. CLSI sensitive/resistant breakpoints.** CLSI breakpoints (µg/mL) for 2016-2017[9] were used to determine antibiotic resistance of clinical isolates.

| Antibiotic | Sensitive | Intermediate | Resistant |
|---|---|---|---|
| Ampicillin (AMP) | 8 | 16 | 32 |
| Ceftriaxone (FRX) | 1 | 2 | 4 |
| Meropenem (MER) | 1 | 2 | 4 |
| Gentamicin (GEN) | 4 | 8 | 16 |
| Kanamycin (KAN) | 16 | 32 | 64 |
| Tetracycline (TET) | 4 | 8 | 16 |
| Ciprofloxacin (CIP) (*E. coli* and *K. pneumoniae*) | 1 | 2 | 4 |
| Ciprofloxacin (*Salmonella enterica*) | 0.06 | 0.125 | 1 |
| Nalidixic Acid (NXA) | 6 | N/A | 32 |
| Chloramphenicol (CHL) | 8 | 16 | 32 |

**Table S2. Unique antibiotic resistance genes identified in clinical isolates.** Italicized label represents antibiotic class the gene confers resistance to where: *Bla* is for β-lactam resistance, *Flq* is for fluoroquinolone resistance, *AGly* is for aminoglycoside resistance, *Phe* is for phenicol resistance, *Tet* is for tetracycline resistance, *Sul* is for sulfonamide resistance, and *Tmt* if for trimethoprim resistance. Non-italicized portion is the unique gene identified by ARG-ANNOT.

| **CRE *E. coli*** | **MDR *E. coli*** | **ESBL KPN** | **NDM-1 KPN** | **STm** |
|---|---|---|---|---|
| *Bla* AmpC1 | *Bla* AmpC2 | *Bla* SHV-11 | *Bla* TEM-217 | *AGly* Aac6-Iaa |
| *Bla* AmpH | *Bla* PBP | *Bla* AmpH | *Bla* NDM-1 | *Bla* PBP |
| *Bla* AmpC2 | *Bla* ampH | *Bla* Oxa-9 | *Bla* CTX-M | |
| *Bla* CMY-94 | *Bla* TEM-219 | *Bla* TEM-171 | *Bla* SHV-73 | |
| *Bla* PBP | *Bla* TEM-10 | *Bla* TEM-220 | *Bla* PBP | |
| *AGly* AadB | *Bla* CTX-M | *Bla* KPC-3 | *Bla* AmpH | |
| *AGly* StrA/B | *AGly* Sat-2A | *Bla* PBP | *Bla* PBP | |
| *Phe* PheCml45 | *Tmt* DfrA1 | *Flq* OqxBgb | *Flq* QnrB1 | |
| *Tet* TetB | *Phe* CatA1 | *AGly* AadA1-pm | *Flq* Qnr-S1 | |
| *Sul* SulI | | *AGly* Aac6-Ib | *Flq* OqxBgb | |
| *Tmt* Dfr24 | | | *AGly* StrB | |
| | | | *AGly* RmtF | |
| | | | *AGly* Ant3 | |
| | | | *Tet* TetA/R | |
| | | | *Sul* SulI | |
| | | | *Tmt* DfrA1 | |

**Table S3. Predicted 0-bp mismatch off-targets of antisense-PNA molecules in each reference bacterial genome and clinical isolate**. The label “STC” denotes gene off-targets for which the antisense-PNA aligns to the start codon, and which will therefore be much more likely to experience translational inhibition.

| **Bacterial Reference Genome Results** | | | |
|---|---|---|---|
| **PNA** | ***E. coli* MG1655** | ***K. pneumoniae* MGH 78578** | ***Salmonella enterica* serovar Typhimurium SL1344** |
| **α-folC** | None | KPN_04193 (putative 6-phosphofructokinase)<br>*uxaC* | None |
| **α-rpsD** | *cueO*<br>*narI* | Non-protein coding region | *wcaM*<br>*rtcA* (STC) |
| **α-ffh** | None | None | None |
| **α-lexA** | None | None | None |
| **α-gyrB** | Last 5 nt of *psiE* | None | None |
| **α-acrA** | *narX* (STC) | MFS transporter (STC), non-protein coding region | None |
| **α-csgD** | Non-protein coding region | Non-protein coding region | Non-protein coding region (2) |
| **α-fnr** | None | None | FNR (non STC) |
| **α-recA** | None | None | None |

| **Clinical Isolates Results** | | | | | |
|---|---|---|---|---|---|
| | **CRE *E. coli*** | **MDR *E. coli*** | **ESBL KPN** | **NDM-1 KPN** | **STm** |
| **α-folC** | Prokka 00542<br>Prokka 03005 | None | *pfka1* | *pfkA1* | None |
| **α-rpsD** | *cueO* | *cueO* | *frlD*<br>Non-protein coding region | Non-protein coding region | Prokka03791<br>Prokka 03488<br>*rtcA* (STC) |
| **α-ffh** | *uhpC* | *uhpC* | *yhes1* | *yheS2* | None |
| **α-lexA** | None | None | None | None | None |

| | | | | | |
|---|---|---|---|---|---|
| **α-gyrB** | Non-protein coding region<br>Last 5 bp of *yhbX* | None | Non-protein coding region | None | None |
| **α-acrA** | *narX* (STC), *livH* | *narX* (STC), *livH* | *exuT* (STC) | *exuT* (STC) | None |
| **α-csgD** | None | Sodium:sulfate symporter | Hypothetical protein | None | None |
| **α-fnr** | None | None | None | None | None |
| **α-recA** | None | None | None | None | None |

**Table S4. DNA oligonucleotides containing PNA-target gene sequence that were utilized for DNA-PNA complex formation in Electrophoretic Mobility Shift Assay.** Synthetic DNA oligonucleotides were 60 nt in length (purchased from IDT), contained the PNA-target gene sequence bolded and underlined. The nonsense oligonucleotide was a randomized 60nt sequence that had no complementarity for any PNA designed for the study served as a non-binding control.

| Oligo/Primer Purpose | Target gene | Oligo/Primer Sequence (5' to 3') |
|---|---|---|
| Antisense oligomer for α-rpsD | *rpsD* | ATT TAG GTG ACA CTA TAG AAG TGG AGA A**AG** **AAA** ***ATG*** **GCA** **A**GA TAT TTG GGT CCT AAG CTC |
| Antisense oligomer for α-lexA | *lexA* | ATT TAG GTG ACA CTA TAG AAG CAG GGG G**CG** **GAA** **TGA** **AAG** **C**GT TAA CGG CCA GGC AAC AAG |
| Nonsense oligomer | N/A | GAA TTC GAA TTC GGT CAG TGC GTC CTG CTG ATG TGC TCA GTA TCT CTA TCA CTG ATA GGG |

**Table S5. Genetic and phenotypic antibiotic resistance characterization.** Pink cells correspond to full resistance across an antibiotic class. Yellow is partial resistance to the class, and green is full sensitivity to the class.

| | **CRE *E. coli*** | **MDR *E. coli*** | **ESBL KPN** | **NDM-1 KPN** | **STm** |
|---|---|---|---|---|---|
| **Antibiotic Type** | **Number of Resistance Genes** | | | | |
| β-lactams | 5 | 6 | 7 | 7 | 1 |
| Aminoglycosides | 2 | 1 | 2 | 3 | 1 |
| Tetracyclines | 1 | 0 | 0 | 1 | 0 |
| Fluorquinolones | 0 | 0 | 1 | 3 | 0 |
| Phenicols | 1 | 1 | 0 | 0 | 0 |
